# Clusterflock: A Flocking Algorithm for Isolating Congruent Phylogenomic Datasets

**DOI:** 10.1101/045773

**Authors:** Apurva Narechania, Richard Baker, Rob DeSalle, Barun Mathema, Sergios-Orestis Kolokotronis, Barry Kreiswirth, Paul J. Planet

## Abstract

**Background:** Collective animal behavior such as the flocking of birds or the shoaling of fish has inspired a class of algorithms designed to optimize distance-based clusters in various applications including document analysis and DNA microarrays. In the flocking model, individual agents respond only to their immediate environment and move according to a few simple rules. After several iterations the agents self-organize and clusters emerge without the need for partitional seeds. In addition to their unsupervised nature, flocking offers several computational advantages including the potential to decrease the number of required comparisons.

**Findings:** In Clusterflock, we implement a flocking algorithm designed to find groups (flocks) of orthologous gene families (OGFs) that share a common evolutionary history. Pairwise distances that measure the phylogenetic incongruence between OGFs guide flock formation. We test this approach on several simulated datasets varying the number of underlying topologies, the proportion of missing data, and evolutionary rates, and show that in datasets containing high levels of missing data and rate heterogeneity, clusterflock outperforms other well-established clustering techniques. We also demonstrate its utility on a known, large-scale recombination event in *Staphylococcus aureus*. By isolating sets of OGFs with divergent phylogenetic signal, we can pinpoint the recombined region without forcing a pre-determined number of groupings or defining a pre-determined incongruence threshold.

**Conclusions:** Clusterflock is an open source tool that can be used to discover horizontally transferred genes, recombining areas of chromosomes, and the phylogenetic “core” of a genome. Though we use it in an evolutionary context, it is generalizable to any clustering problem. Users can write extensions to calculate any distance metric on the unit interval and use these distances to flock any type of data.

## Findings

### Background

Swarm intelligence describes the cooperative behavior that results from a group of agents executing simple behavioral programs. The agents themselves are unsophisticated, but patterns emerge from the accumulation of pairwise interactions resulting in completion of complex tasks necessary for the group’s survival [1]. Swarms are by definition leaderless and the agents are given no internal or external direction. Ant colonies, swarms of bees, shoals of fish, and flocks of birds all demonstrate this kind of behavior [2-5].

Because there is no central control, algorithms modeled on swarm intelligence excel at data mining tasks where the goal is discovery of unknown patterns in data. Early models of this type of behavior invoked the N-body problem as studied in the physics of celestial objects [6]. Later work integrated biological observations, particularly the importance of local information in determining global patterns of behavior [7-10]. The Reynolds flocking model [11] is one such example designed specifically to simulate the coherent behaviors characteristic of a flock of birds. Reynolds original goal was to bestow life-like animation to particles – which he termed *boids* – in motion pictures. In Reynolds’ flocking, each boid in a simulation is a clone of every other, and boids heed only their immediate surroundings as delimited by a radius of perception. Boids react to flockmates within this radius using a small library of simple behaviors that ultimately result in the synchrony of the entire group. If boids are assigned bits of information and if distances between these bits of information are easily computed, then the flocking algorithm becomes suitable for unsupervised clustering.

Here we present Clusterflock, a method that aims to isolate groups (flocks) of genes with congruent historical signals. In our technique, boids represent orthologous gene families (OGFs), and the distance between them is measured as a simple test of phylogenetic congruence [12].

Phylogenetic incongruence is rampant in the evolutionary history of genes across most organisms in the tree of life [13-16], but the problem is particularly severe among bacterial genomes where evolution proceeds through multiple mechanisms that destroy phylogenetic signal including recombination, *de novo* gene acquisition, loss and duplication [17]. A central question in microbial evolutionary biology is how to distinguish these non-vertical mechanisms and separate vertical signal from the horizontal signal produced by recombination and gene transfer. Whole genomes have thousands of genes with potentially different histories magnifying the analytical and computational complexity of this problem [18, 19]. The number of recombination events and the rates of gene transfer are often not known, and inclusion of genes with different histories in the same analysis can lead to bias and errors in phylogenetic trees. This kind of error is especially problematic in cases of large-scale recombination or sustained/concerted gene transfer between organisms that occupy the same habitat [20], situations that can lead to strong support for the incorrect hypothesis.

A handful of algorithms have been put forward to address this phylogenetic problem in large whole-genome datasets. In general, such approaches rely on a two-step procedure: pairwise tests of phylogenetic incongruence between OGFs followed by clustering to segregate OGFs into congruent groups. A range of initial incongruence tests have been used including character-based incongruence measures such as the incongruence length difference [e.g.,mILD [19]] or the likelihood ratio test [e.g., CONCATERPILLAR [21]], and topological measures [e.g., Conclustador [18]]. For the clustering step, CONCATERPILLAR and mILD use agglomerative, hierarchical clustering techniques, whereas Conclustador uses k-means and spectral clustering algorithms. Hierarchical clustering techniques generally require a threshold value that defines the boundaries of groups, an assumption that could introduce bias or error. Spectral, as implemented in conclustador, and k-means algorithms require prior estimation or specification of the number of clusters, which could also lead to erroneous lumping or splitting. In contrast, Clusterflock does not require prior specification of distance thresholds or the number of groups.

We have used the incongruence length difference (ILD) in our analysis of the Clusterflock algorithm, but the algorithm could also be applied to the likelihood ratio test or the topological incongruence measure presented in Conclustador [18]. Indeed the flocking algorithm can cluster anything given precalculated pairwise distances between all entities. Here we test the flocking model against other clustering algorithms such as multidimensional scaling (MDS), hierarchical clustering, and partitioning around medoids

(PAM). To demonstrate its utility in the real world, we also use Clusterflock to analyze a well-studied example of massive genome recombination in *Staphylococcus aureus* clonal group ST239 involving nearly 20% of the chromosome [22]. Like many phylogenomic datasets, the staphylococcal genomes we analyzed have large amounts of missing data, a circumstance that often limits the effectiveness of clustering algorithms [18]. While other techniques failed to segregate the recombined region of the ST239 genome, Clusterflock successfully distinguishes recombined genes from those in the recipient genome. We have explored this resilience to missing data and to rate heterogeneity through simulation.

### Implementation

A depiction of Reynolds’ original algorithm and our modifications is shown in Figure 1. At the outset, agents (boids in the simulation) are assigned a random position and velocity in a 2-dimensional field normally set at one particle per square unit. Agents are then allowed to interact with one another. In the classic approach, individual entities are influenced only by their local environment as given by a user-defined radius. For each agent, flockmates within this radius will influence the calculation of three steering vectors that combine to alter the particle’s velocity: *Alignment, Cohesion* and *Separation*. Agents will tend to head in the average direction of their flockmates (alignment), move towards their average position (cohesion), and avoid crowding one another (separation). To accelerate the formation of flocks, we added *Repulsion* as a fourth vector and designed it to operate between each agent’s field of vision (radius) and its radius of separation. In a clustering context, repulsion quickly separates agents with high relative distances, seeding flocks at an early point in the simulation. The calculation of the four vectors is iterated over all agents in the system for a user-defined number of frames.

**Figure 1.**
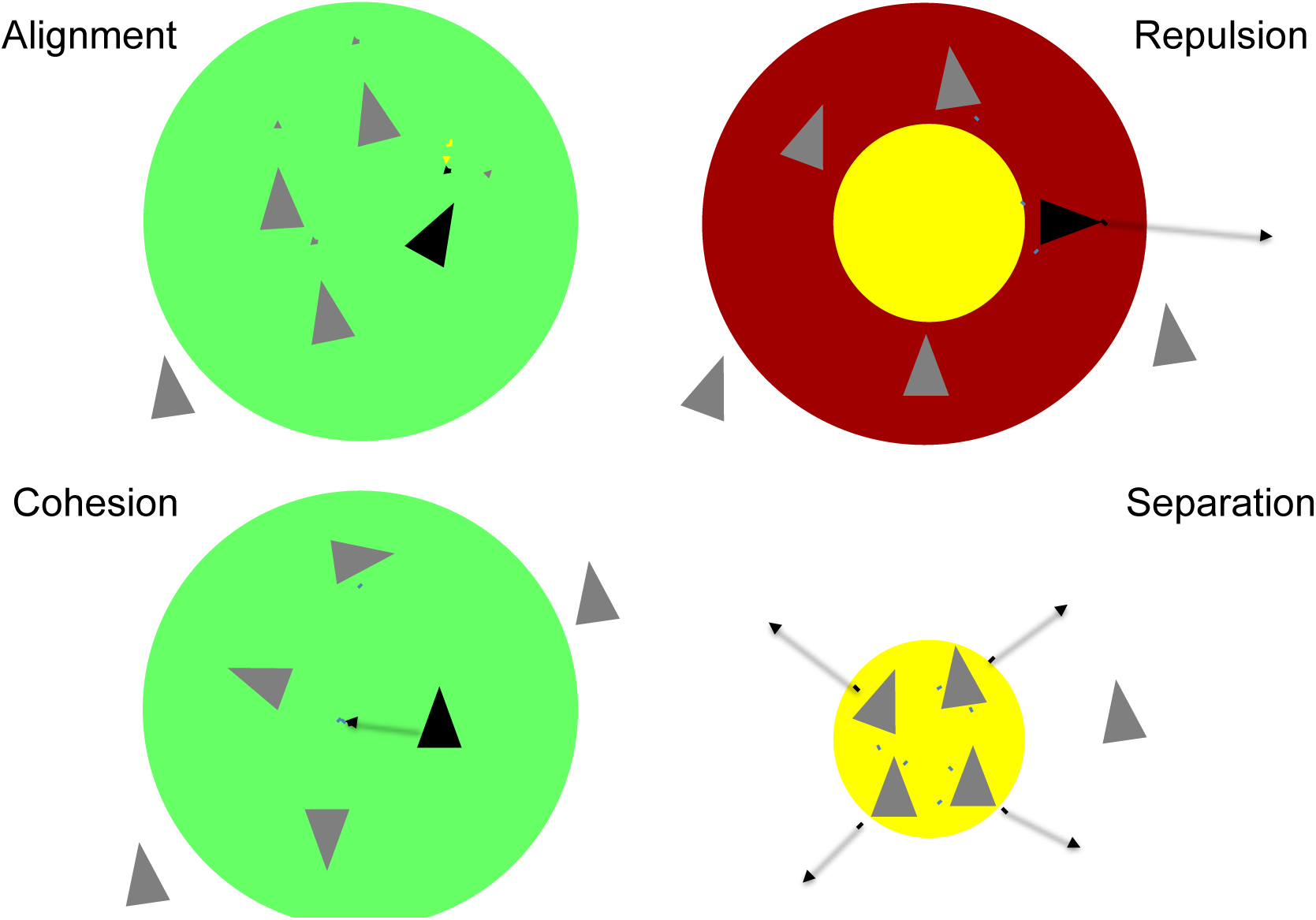
Flocking Algorithm and Rules. Individual agents (boids) are shown as triangles, interactions as dashed lines, and the radius of perception as differently colored circles depending on the vector considered. Alignment and cohesion reinforce flocking behavior while repulsion disrupts it. In cohesion, a given boid moves towards the center of mass of all congruent flockmates within its field of vision. Boids will also align their velocities with congruent flockmates. However, all boids regardless of whether they are congruent or incongruent will separate from flockmates in their immediate vicinity. Cohesion, alignment, and separation are the core forces in Reynold’s original flocking algorithm. We have added repulsion as a force capable of isolating congruent and incongruent agents. It operates between an agent’s field of vision and the smaller, concentric circle describing this agent’s radius of separation. The magnitudes of alignment, cohesion, and repulsion are a function of the phylogenetic distance between the agents as described in the implementation.

In our adaptation, each particle is a set of OGFs in sequence alignment (i.e., a phylogenetic matrix), and each interaction triggers a measurement (or a hash table lookup) of phylogenetic congruence between alignments in the pair. We model our congruence metric after the incongruence length difference (ILD) [23]:

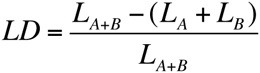

where *L*_*A*+*B*_ is the length of the Maximum Parsimony (MP) tree calculated when the two gene alignments are combined (i.e. concatenated), and *L*_*A*_ and *L*_*B*_ are the lengths of the trees calculated for each gene individually. An LD of zero indicates complete congruence (i.e. the gene trees are identical), whereas a positive LD indicates that the two OGFs have divergent phylogenetic topologies. To function as a distance metric on the unit interval, LD is normalized and therefore scaled between 0 and 1. We calculated single gene parsimony trees and concatenated trees in PAUP* [24] using 100 heuristic searches with random sequence addition and tree bisection and reconnection.

Unlike other implementations [25, 26] we use the LD metric directly in our formulation of the steering vectors. This allows the distance between any two OGFs to have a continuous effect on the simulation. More specifically, alignment and cohesion are calculated as the average velocity and position, respectively, of all flockmates within any given agent’s field of view (Figure 1). This average is then modulated by two factors: the average LD of all flockmates, and a user-defined diminishment factor. The equation for alignment is shown here.

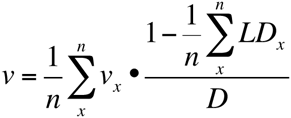

where *v* is the velocity driven by alignment, *n* is the number of flockmates, *v*_*x*_ is the velocity of flockmate *x*, *LD*_*x*_ is the LD for flockmate *x*, and *D* is the diminishment factor.

In this scenario, as the average LD increases, the alignment or cohesion effect decreases. Similarly, as the diminishment factor increases, the alignment or cohesion effect decreases. The diminishment factor is intended as a layer of control, giving the user the opportunity to up weight or down weight the alignment and/or cohesion vectors.

Separation and repulsion are treated somewhat differently and are instead calculated through iterative displacement. Each agent within the separation distance updates the separation vector in turn, attempting to double the distance between itself and its counterpart:

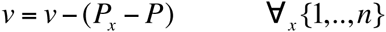

where *v* is the velocity driven by separation, and *P*_*x*_ and *P* are the positions of the agent in question and its flockmate *x*, respectively. Repulsion is enhanced separation that operates between the separation distance and the perception radius. It is directly proportional to the LD of flockmate X and a user-defined enhancement factor:

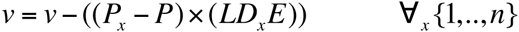

where *LD*_*x*_ is the LD for flockmate *x*, and *E* is the enhancement factor. In this formulation, if the LD between any two OGFs is zero, the repulsion effect vanishes. For positive LDs, repulsion is positive and proportional to the magnitude of the incongruence calculated. Increasing the enhancement factor can magnify any existing repulsion considerably.

Summing the velocities derived from the cohesion, alignment and separation/repulsion rules encodes the evolutionary distance information between an active agent and all its flockmates. Done across all agents over all iterations, OGFs converge on multiple evolutionary solutions, and the final frame of the simulation often isolates all congruent clusters. Figure 2 captures snapshots of this process. Seed clusters form very early and later move to intercept one another. Congruent flocks will absorb one another while incongruent flocks repel.

**Figure 2.**
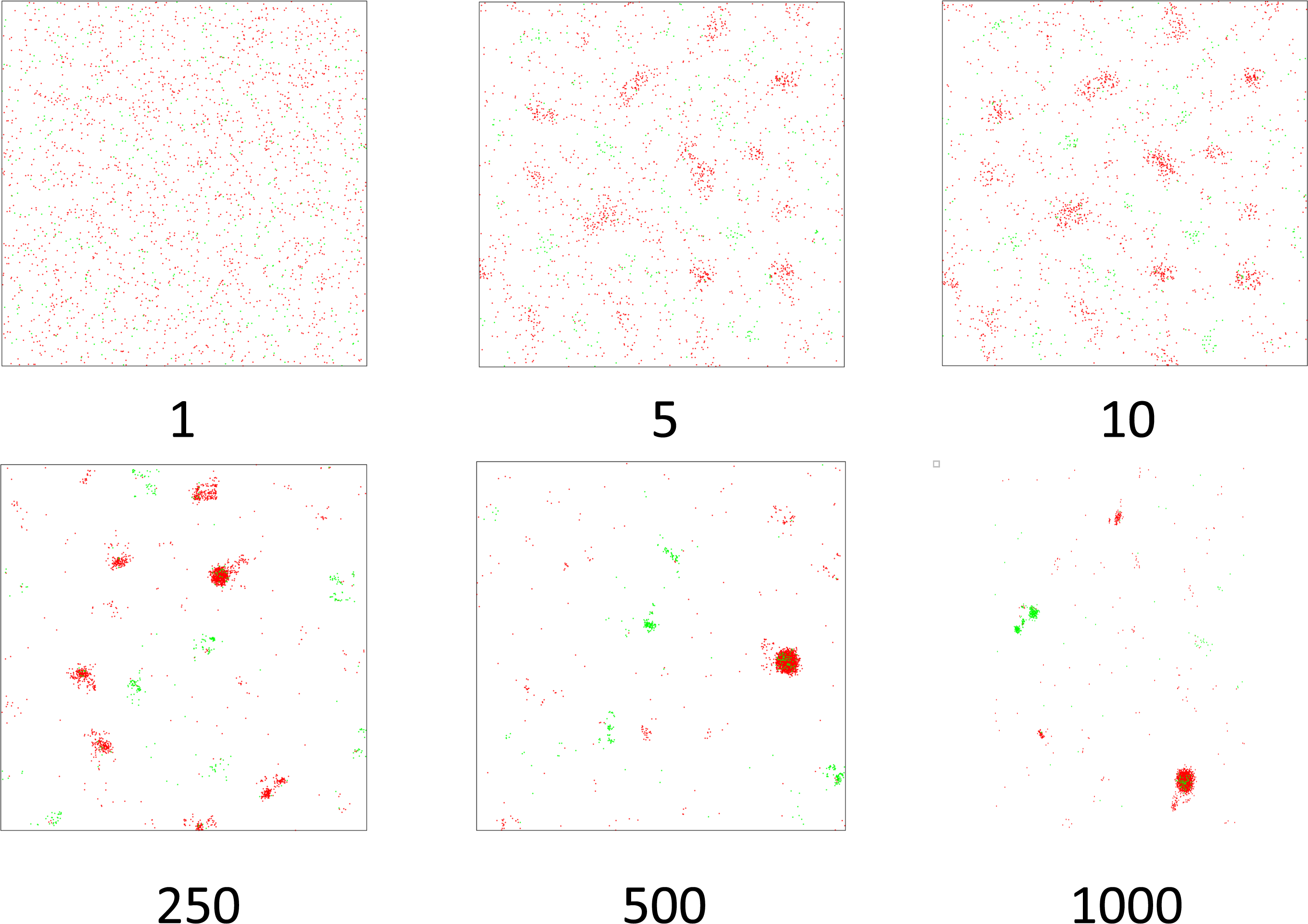
Snapshots from a *Staphylococcus aureus* Simulation. Here we show three early snapshots, and three snapshots taken from intervals along a 1000 frame flocking simulation. Agents colored green represent genes from the recombined region, while those colored red are from the core genome. Specific parameters chosen here include: DIMMENSIONS = 2500; BOUNDARY = 1; INIT_VELOCITY = 50; COHESION_FACTOR = 5; SEPARATION_DISTANCE = 5; REPEL_FACTOR = 10; ALIGNMENT_FACTOR = 5; ITERATIONS = 1000; RADIUS = 500; VELOCITY_LIMIT = 50; MINPTS=20; XI=0.15.

Clusterflock is a parameter-rich approach, allowing the user fine-grain control over the steering of OGFs within the virtual space. Besides the cohesion, alignment and repulsion factors outlined here, other key parameters include the length of the virtual square that serves as the flight space, the radius of awareness around each agent, the initial velocity, the velocity limit, and the number of iterations. We have found that a gene per square unit and a radial awareness of 10% of the flight space length are sufficient to encourage efficient flock formation in most cases. Since velocity can increase quickly if left uncapped, limiting it to 2% of the virtual square’s length is usually adequate for circulation.

In its original form, the algorithmic complexity of flocking is *O*(*n*^2^). Each agent must spatially assess all other agents to determine who is in its field of view (radius). We have reduced this complexity by borrowing two heuristics, one from the world of video games, and one from nature itself. In video games that require real-time calculation of the interactions of many particles, a spatial hashing structure [27, 28] reduces the number of required comparisons by binning particles into a discrete number of cells. Agents are sorted by their location and only those in cells immediately surrounding the query are processed. In practice, spatial hashing’s most significant savings are accrued early in a simulation when agents are evenly dispersed. As flocks begin to form, some cells are completely bereft, while others can contain thousands of congruent OGFs.

A more even and lasting heuristic is awareness. In nature, an individual in a flock will not necessarily respond to *all* its immediate flockmates [10, 29]. Often, it responds to a mere subset, an approximation that rarely leads to unwanted perturbations or collisions because of the cumulative, emergent nature of the group’s motion as a whole. Capped by a user-input maximum awareness, a random sampling of flockmates for each gene is often sufficient to guide groups of flocks into distinct evolutionary classes. This heuristic can be extremely significant towards the end of a simulation when many of the final flocks have formed. Groups of thousands of congruent individuals are not uncommon.

By dimming each agent’s effective perception, only a fraction of these congruent individuals require analysis.

Despite these shortcuts, flocking is a computational expensive procedure. Without image/movie creation and analysis of cluster formation progress (k-means or OPTICS, see below), the flocking procedure alone will require on average 30 CPU seconds for a dataset containing 100 loci of 100 residues each across 10 taxa. Because we encourage 100 or more replicates, total required CPU per experiment for this hypothetical dataset would be just short of 1 CPU hour. In contrast, a single run of other clustering procedures like MDS, hierarchical clustering, and PAM require less than a CPU second to analyze this type of data. For clusterflock, what is lost in terms of speed is gained in performance as we show in the next section.

### In test: simulations and comparisons to dominant clustering techniques

We simulated 100 locus datasets containing 100 residues per locus across 10 taxa in Seq-Gen [30] using JTT. Underlying these simulated proteins were anywhere from 1 to 25 generated topologies: in the case of 1 underlying topology all 100 loci were modeled as congruent; in the case of 25 topologies loci were randomly assigned to each tree without requiring that each tree be equally represented. Because missing data is a common problem in phylogenomics we also chose to model data sparsity’s effect on clustering performance. Taxa were randomly assigned as missing on a per locus basis at rates varying from 0% to 50% in 10% increments. Because rate heterogeneity is also a common problem, in a separate set of experiments with no missing data, we randomly assigned about half the proteins in each matrix to be 3X (1.5/0.5), 7X (1.75/0.25) or 19X (1.9/0.1) faster than their counterparts in terms of relative evolutionary rates [30]. For statistical power we repeated dataset creation 10 times per topological condition across our missing data thresholds and rate heterogeneity multipliers resulting in a grand total of 2250 matrices.

In addition to using clusterflock (100 replicates of 500 frames each), we analyzed the 100 proteins in each matrix using multidimensional scaling (MDS), hierarchical clustering and PAM, using the R packages [31] cmdscale, hclust, and pam, respectively. In the case of PAM, a k-medoids operation, and hierarchical clustering users must provide an estimate of expected cluster number at the outset. In practice, providing this number is often difficult given the increasing complexity of modern phylogenomic datasets. We instead favor Clusterflock and MDS, techniques that spatially encode the distance information between loci allowing for the emergence of distinct data categories.

Figure 3 summarizes the results of these simulations. Shown is the average Jaccard Index of all replicates against the number of simulated topologies across all data sparsity thresholds. Because we know which tree is associated with each gene, we can employ the Jaccard Index as an external measurement of clustering efficiency. The Jaccard is defined as follows:

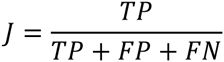

where TP is true positive, FP is false positive and FN is false negative as judged by correct assignation to a congruent tree topology group.

**Figure 3.**
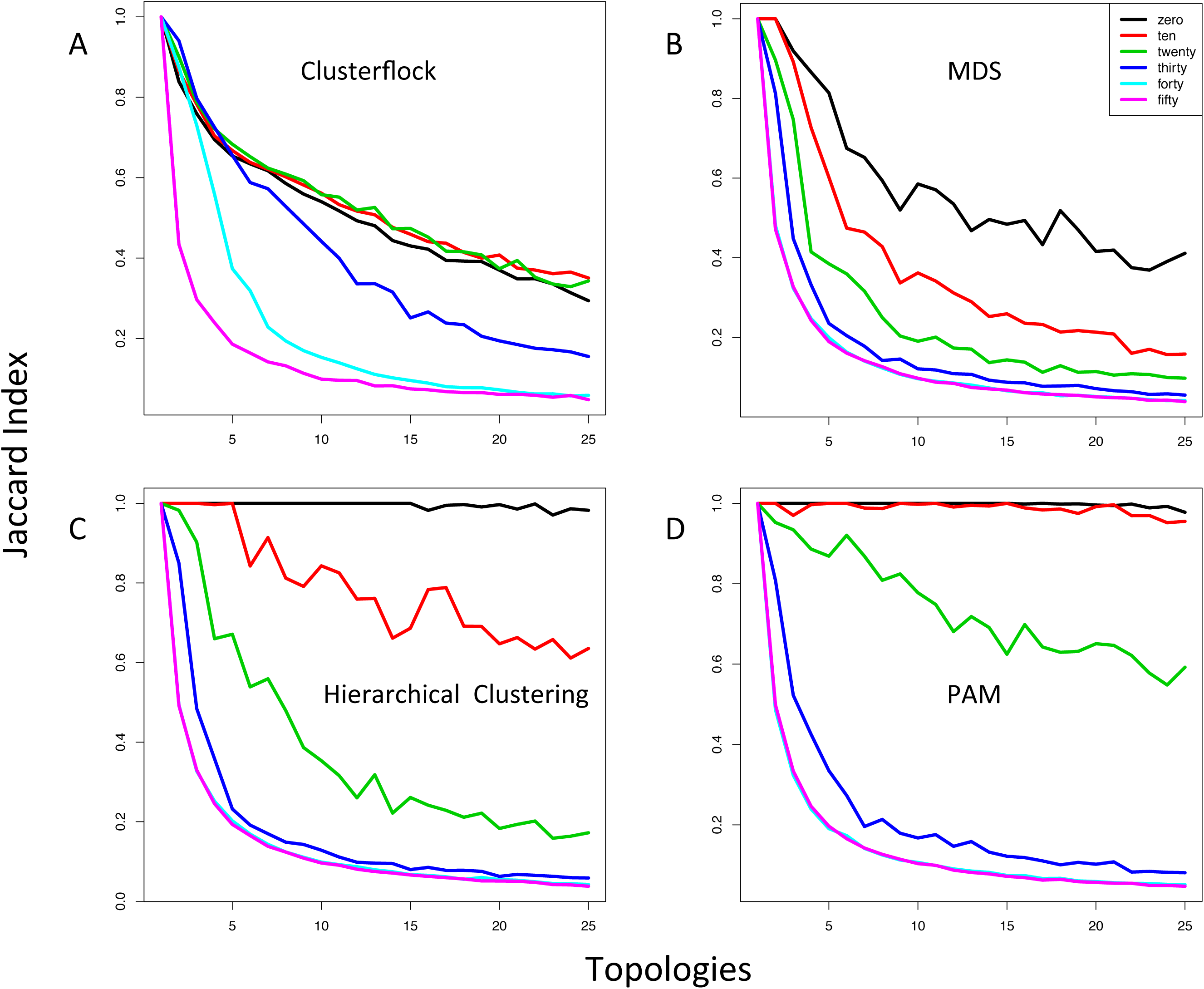
Simulating Topological Complexity and Missing Data. Four plots measuring the Jaccard Index (see In test) against increasing topological complexity and across increasing levels of missing data percentage (colored as indicated in the insert) are shown. We compare four methods here: (A) Clusterflock, (B) multidimensional scaling, (C) hierarchical clustering, and (D) Partitioning around medoids (PAM). In order to compare the four techniques using similar methods, and because the number of underlying topologies in each simulation is known, for Clusterflock and MDS we used k-means to cluster the final spatial arrangement of loci and assign OGFs to topological groups.

As expected, the performance of all four methods degrades with increasing topological complexity and increasing data sparsity. No method performs well when 50% of the data is missing. And increasing the topological heterogeneity makes it more difficult for any method to discern between groups. However, Clusterflock is robust to the effects of missing data in a way that no other method tested here has shown. It also mirrors the performance of MDS as the evolution of the underlying genes becomes more complicated.

With the advantage of seeding, hierarchical clustering and PAM,are superior in the special case where the number of topologies are absolutely known and that data is used to guide cluster formation. As long as there is little missing data, these two methods are clearly superior among the techniques we compare here. Surprisingly, hierarchical clustering and PAM perform very poorly as data is removed. For MDS and Clusterflock, methods that spare the user an initial estimation of the number of clusters, increasing topological complexity lowers the Jaccard at approximately the same rate as long as there is no missing data. The difference between MDS and Clusterflock techniques materializes once the data thins.

Figure 4 shows the effect of rate heterogeneity. MDS degrades rapidly with increasing rates of relative heterogeneity, while the strong performance of Clusterflock persists across evolutionary rates. Therefore, up to a point, clusterflock is resilient to both missing data and differing evolutionary rates. Zero, ten and twenty percent missing data perform equally well whereas MDS shows quick collapse as information is removed. And diverging evolutionary rates among genes do not seem to affect clusterflock’s ability to sort them into their respective topological groups.

**Figure 4.**
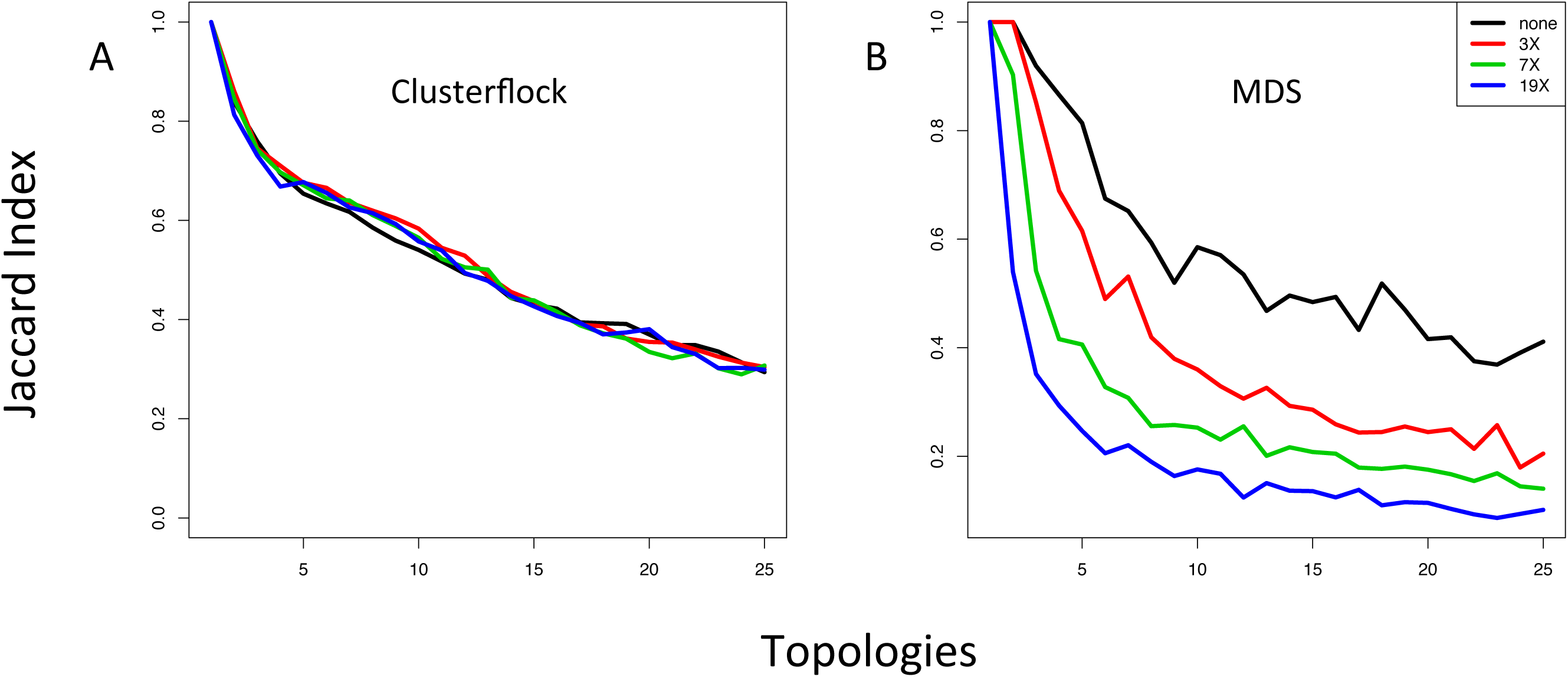
Simulating Topological Complexity and Evolutionary Rate Heterogeneity. Here we show two plots measuring the Jaccard Index against topological complexity and across four levels of relative evolutionary rate: 1X (1/1); 3X (1.5/0.5); 7X (1.75/0.25) and 19X (1.9/0.1) (colored as indicated in the insert). We compare two methods here: (A) Clusterflock, and (B) multidimensional scaling.

### In action: a recombination event in Staphylococcus aureus

To test our ability to detect and separate distinct populations of incongruent OGFs in biological systems, we used a well-known example of large-scale genomic recombination between two S. *aureus* clonal complexes (CC) [22]. The genomes of ST239 S. *aureus* appear to have formed from a recombination event in which 20% of a CC8 genome was replaced with the homologous portion of the genome from a CC30 strain. We chose 11 S. *aureus* strains (GCA_000146385.1, GCA_000012045.1, GCA_000011505.1, GCA_000011265.1, GCA_000013425.1, GCA_000204665.1, GCA_000159535.2, GCA_000027045.1, GCA_000017085.1, GCA_000236925.1 and SA21300) including examples from CC30, CC8 and ST239 and generated groups of orthologous genes across all their proteomes using orthologID [32]. The resulting sequence data matrix contained 2550 orthologous OGFs totaling 758,270 amino acid characters. Missing data was tolerated, but representation from at least 4 taxa for each gene was required.

Figure 2 shows three frames from the beginning of a 1000 frame simulation and three snapshots from even intervals thereafter. We have color-coded OGFs here for clarity: green maps to the hybridized portion of the genome, and red maps to the remainder. A complete video of this simulation can be found at https://youtu.be/v_4bDprmkpU.

Sorting of OGFs by phylogeny is evident as early as the 10^th^ frame and proceeds further as these initial seeds encounter one another in the virtual space. But hundreds to thousands of frames are required to amass flocks containing all congruent OGFs. The number of frames required is dependent on the general phylogenetic cohesion of the OGFs, and their random starting positions and velocities in the virtual space. By the end of the simulation, the OGFs have self-organized in the leaderless way characteristic of swarm behavior. Without having to estimate the number of expected evolutionary trajectories, we find that there are two dominant flocks: one corresponding to the recombined region and the other to the rest of the genome. We observe two unexpected, smaller flocks (Figure 5) whose provenance does not trace to any known evolutionary event or functional class.

**Figure 5.**
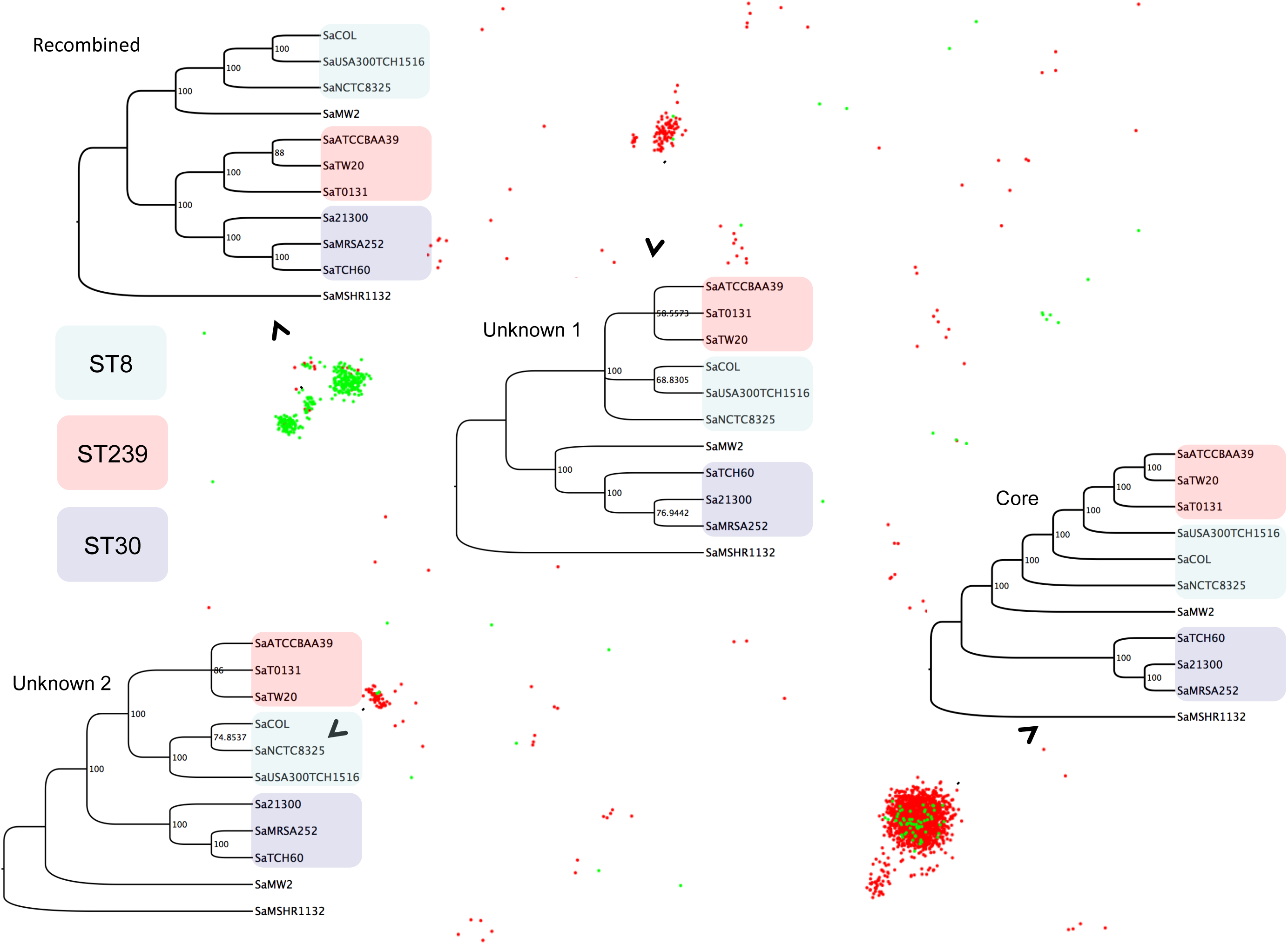
The Final Frame. Four parsimony trees corresponding to the four dominant flocks are shown superimposed on the final frame of the example simulation. Taxa are colored according to their phylogenetic group (MLST classification). Genes from the recombined region place ST239, the hybrid strains, sister to ST30, whereas genes from the rest of the genome yield a monophyletic arrangement of ST239 and ST8. The unknown phylogenies highlight genes that are members of two novel evolutionary histories.

Because of the stochasticity inherent in this type of behavioral method there is no guarantee that flocks of the recombined region or the rest of the genome will be complete. Either or both may enter the final frame in pieces. In other words, we need more statistical heft than one simulation can provide. To test the reproducibility of the flocks and identify robust flock membership, we repeat the simulation in parallel, randomly varying each OGF’s initial placement and velocity. For the purposes of this example, we deployed 100 replicates.

Visual inspection of 100 final frames, or the thousands that may be desirable for other clustering problems, is prohibitive. We automate the analysis of each final frame using ELKI’s [33] implementation of the OPTICS algorithm [34], a method designed to identify unseeded clusters in spatial data regardless of their density. Note that in contrast to our simulations where we know the number of clusters and can therefore leverage k-means, here we assume that this number is unknown, and in the case of very complex datasets, unknowable. Supplemental Figure 1 shows the average number of auto-detected flocks across all 100 simulations as a function of their frame. After an initial spike, or seeding stage, micro-clusters combine with neighboring micro-clusters that share a congruent phylogenetic history. A near exponential decay in the number of auto-detected flocks is followed by a steady state. In the case of S. *aureus*, by the 200^th^ frame, Clusterflock has isolated most of the evolutionary paths.

We systematize the flocking information by assigning a flock label to each OGF for each replicate. For example, replicate 1 may have resulted in five flocks; each of its 2550 gene families is therefore assigned to A, B, C, D, or E. Similar assignments are made across all replicates and the labels are treated as character information in a matrix. The Flock Matrix therefore has as many rows as there are OGFs and as many columns as there are replicates. Organized in this way, we can derive our final sets of congruent OGFs using tree reconstruction as shown in Figure 6A. This topology highlights groups of OGFs that flock together regularly across all replicates. Here we employ Neighbor Joining and assess node robustness by bootstrapping 100 times. Since we have no external truth against which to measure success, we propose that the application of non-parametric bootstrapping to the Flock Matrix can serve as the basis for assessing validity [35-38]. We observe high levels of support for the flock that is composed of recombined genes, and flocks that represent the two novel phylogenetic histories highlighted in Figure 5.

**Figure 6.**
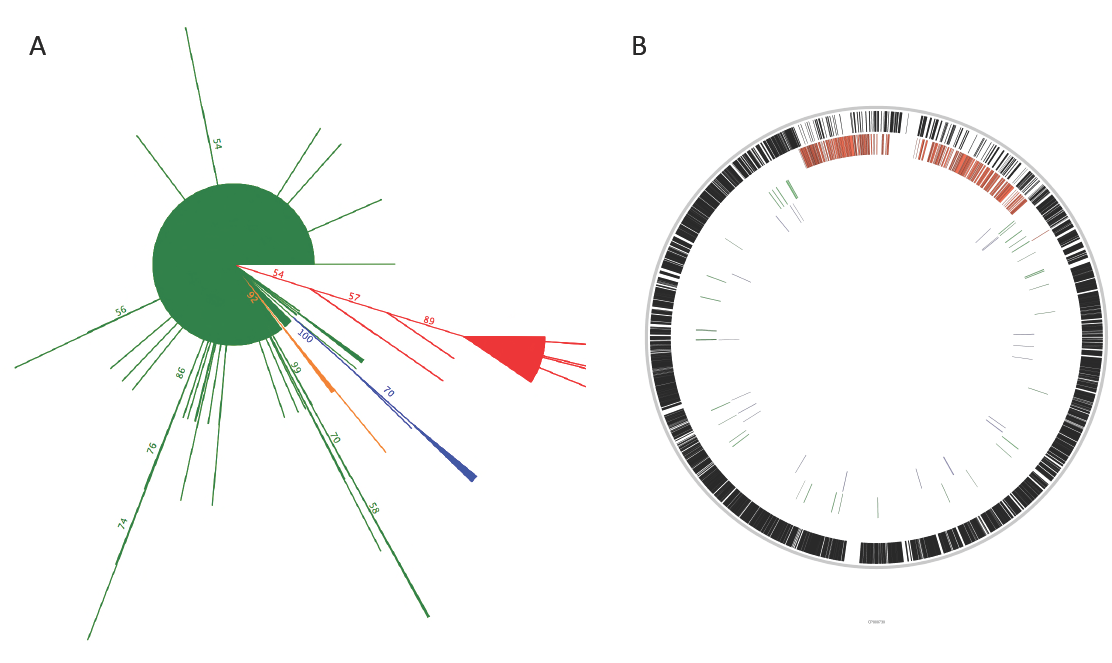
Consensus tree of *Staphylococcus aureus* Flocks Mapped to the USA300TCH1516 Genome. (A) The neighbor joining bootstrap consensus tree for 100 simulations is shown. The majority of the genome occupies the largest branch comprising a virtual polytomy (red). Four other branches of note are highlighted, the largest of which describes the flock consisting of genes from the recombined region (green). (B) We constructed HMMs from the orthologous groups (genes) in each of the four dominant flocks and queried them against a USA300TCH1516 reference. Their genomic locations are shown in the four tracks displayed here. The outermost track is composed of genes from the largest flock. The second track localizes genes from the second largest flock to the known recombined region.

Our main premise was that we could use Clusterflock to detect the genes involved in the ST239 recombination event *a priori* with just pairwise interactions guided by incongruence metrics as OGFs encountered one another in a virtual space. There was no expectation that we would find only two unique flocks, an assumption required by other clustering methods keyed on partitional seeds. Indeed, two additional flocks emerge, with OGFs that tightly share a unique historical signal distinct from our two main evolutionary classes. When we cluster these data with MDS (Supplemental Figure 2) there is no discernable pattern, a predicament most likely due to some combination of missing data and rate heterogeneity.

If, as we suspect, the conflicting histories of our two main flocks originate from the recombination event, we should see the gene families sort based on the known boundaries of the structural change. By creating profile HMMs [39, 40] of each of our 2550 OGFs and mapping them to the S. *aureus* USA300/TCH1516 genome using hmmsearch in the HMMER package, we show that the flock of OGFs corresponding to the recombined region (Figure 6B), map there almost exclusively. The flock corresponding to the rest of the genome is enriched in genes outside of the recombined region, but this enrichment is imperfect. Many genes from within the recombined region contaminate the flock representing the rest of the genome. The length differences of these genes with respect to genes in the largest flock is zero, indicating that they are 100% congruent with the non-recombined phylogeny. These select regions of the recombination event could have reverted through a series of subsequent recombination events, or may show the original recombination event did not replace one continuous section of the chromosome. A discontinuous pattern of recombination is known to occur in other bacteria [41, 42]. Other possibilities include convergent changes, or high levels of conservation prior to the recombination event.

### Conclusions

Clusterflock is an enhanced version of Reynolds’ original flocking algorithm customized to function as a clustering technology. Here we show that it is well suited to isolating congruent gene families into discrete flocks even if they have significant levels of missing data or rate heterogeneity. It can be used to identify a phylogenetic core of genes that share a vertical evolutionary signal while highlighting genes that conflict in subtle ways. However the technique is general, and not restricted to evolutionary analysis [25]. Any distance metric scaled between 0 and 1 can be used to cluster any set of entities. In an era when supervised machine learning often captures many bioinformatic headlines, Clusterflock is in the tradition of data mining: a bio-inspired clustering algorithm used to discover categories of entities without any training, any sense of the number of categories to expect, or any bias in how distant two entities must be to be considered different.

**Supplemental Figure 1.**
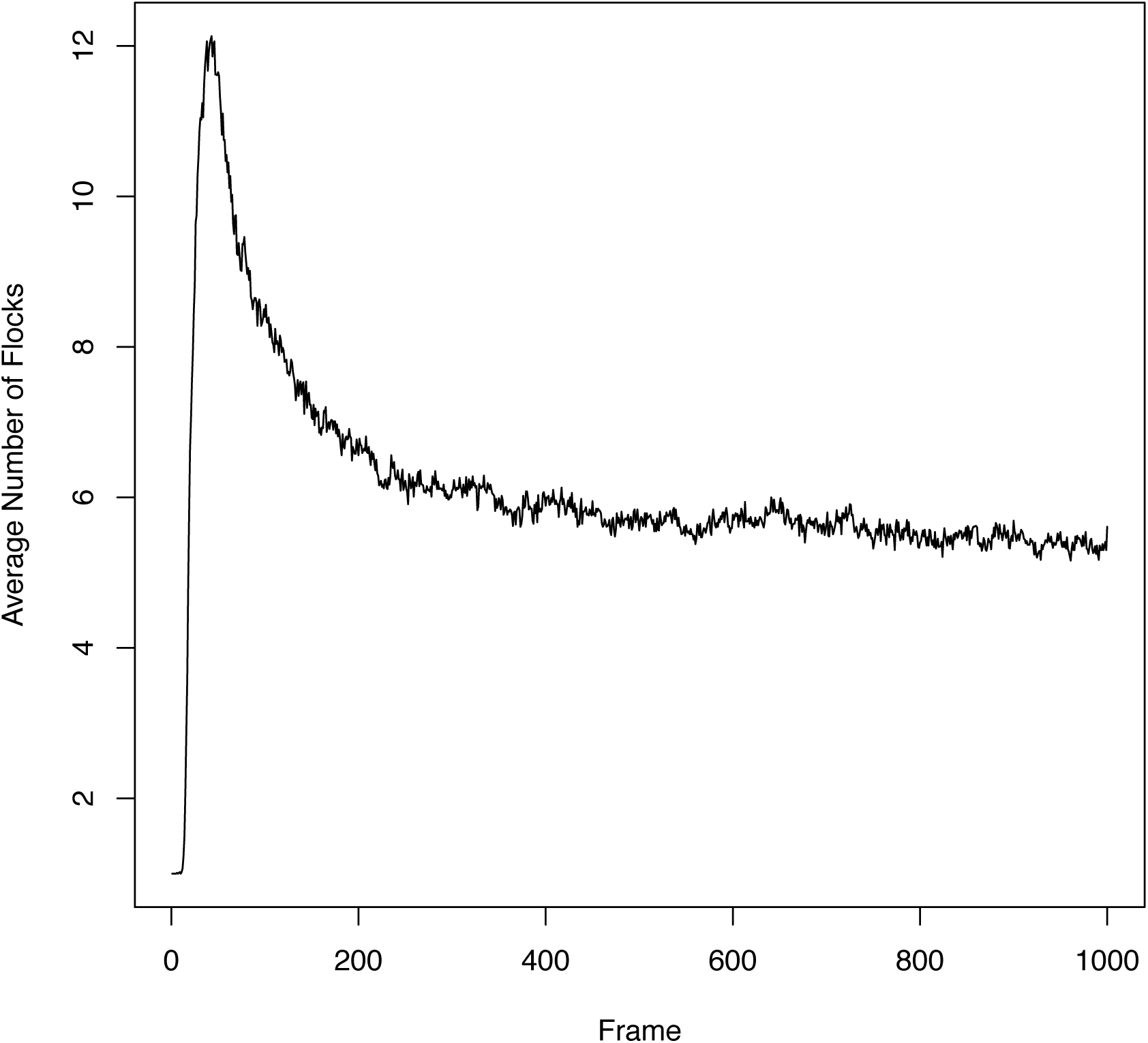
Auto-detected Flocks per Frame. Here we show the average number of flocks detected at any given point along a 1000 frame simulation for the *S. aureus* simulation. The OPTICS spatial clustering algorithm was used to auto-detect flocks in the 100 replicate frames at each point along simulation.

**Supplemental Figure 2.**
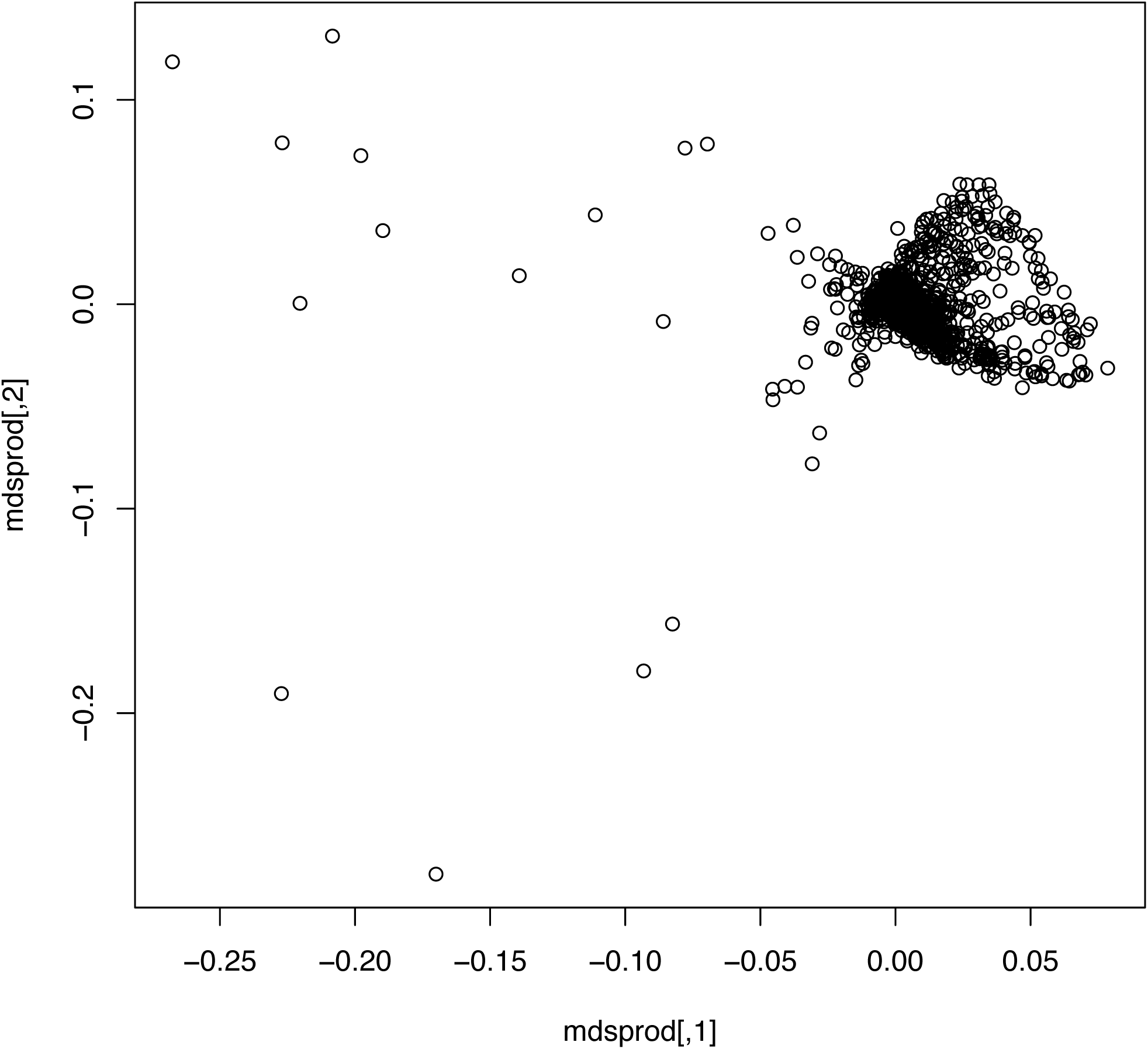
Multidimensional scaling of the *Staphylococcus aureus* dataset. We show a failed attempt to cluster the *Staphylococcus aureus* dataset with MDS. Most loci gather in a single area and we see no separation between the recombined region and the rest of the genome.

## Availability and Requirements

Project Name: Clusterflock

Project Page: https://github.com/narechan/clusterflock

Operating System: Linux

Programming Language: PERL

Other Requirements: See manual in the distribution

License: GPLv3

## Availability of Supporting Data

The data sets supporting the results of this article are available in the GitHub repository. See https://github.com/narechan/clusterflock/tree/master/releases/0.1/example_data for the S. *aureus* data. See https://github.com/narechan/clusterflock/tree/master/releases/0.1/test_data for test data.

